# Environmental unpredictability shapes glucocorticoid regulation across populations of tree swallows

**DOI:** 10.1101/2019.12.23.887075

**Authors:** Cedric Zimmer, Conor C. Taff, Daniel R. Ardia, Alexandra P. Rose, David A. Aborn, L. Scott Johnson, Maren N. Vitousek

**Affiliations:** Department of Ecology and Evolutionary Biology, Cornell University, Ithaca, NY 14853, USA; Cornell Lab of Ornithology, Ithaca, NY 14850, USA; Department of Biology, Franklin and Marshall College, Lancaster, PA 17604, USA; Institute of Arctic and Alpine Research, University of Colorado, Boulder, CO 80303, USA; Biology, Geology and Environmental Science, The University of Tennessee Chattanooga, Chattanooga, TN 37403, USA; Department of Biological Sciences, Towson University, Towson, MD 21252, USA

## Abstract

Responding appropriately to challenges is an important contributor to fitness. Variation in the regulation of glucocorticoid hormones, which mediate the phenotypic response to challenges, can therefore influence the ability to persist in a given environment. We compared stress responsiveness in four populations of tree swallows (*Tachycineta bicolor*) along an environmental gradient to evaluate support for different selective pressures in driving the evolution of glucocorticoid regulation. In accordance with the environmental unpredictability hypothesis, stronger stress responses were seen in more unpredictable environments. Contrary to the reproductive value hypothesis, the stress response was not lower in populations engaging in more valuable reproductive attempts. Populations with stronger stress responses also had stronger negative feedback, which supports a “mitigating” rather than a “magnifying” effect of negative feedback on stress responses. These results suggest that combining a robust stress response with strong negative feedback may be important for persisting in unpredictable or rapidly changing environments.

## Introduction

Global environmental changes are altering the habitats of many species (Peñuelas et al., 2013). For species with wide geographic ranges, intra-specific variation in life-history strategies resulting from historical selection might predispose some populations to be more or less susceptible to increasing environmental changes (Candolin, 2019; Debecker and Stoks, 2019; Harding et al., 2019). Environmental variation favors individuals that differentially allocate time and energy to reproduction and self-maintenance in order to maximize lifetime fitness (Stearns, 1989; Stearns, 1992; Ricklefs and Wikelski, 2002). Thus, characterizing differences in the regulation of this trade-off across environments is critical for understanding the mechanisms that have shaped phenotypic responses and that may allow successful adaptation and population persistence under rapid global changes. While the ultimate reasons for variation in life-history traits across environments and latitudes have been well studied, we have a limited understanding of the proximate mechanisms that underlie this variation (Eikenaar et al. 2012, Atwell et al. 2014). Because environmental factors may influence the evolution of life-history traits by acting on physiological systems that integrate external conditions, hormones have been proposed to play a crucial role as mediators of life-history trade-offs (Zera and Harshman, 2001; Ricklefs and Wikelski, 2002; Eikenaar et al., 2012).

The hypothalamic-pituitary-adrenal (HPA) axis is a fundamental component of the endocrine system that forms an interface between an animal and its environment (Wingfield and Sapolsky, 2003; Breuner, 2011; Hau et al., 2016). The HPA axis coordinates the response to energetic and other challenges mainly by regulating the production and release of glucocorticoids (Wingfield et al., 1998; Sapolsky et al., 2000). In non-chronically or acutely stressed individuals, glucocorticoids are usually maintained at low levels to regulate energy balance and mediate foraging and other locomotor activities (Sapolsky et al., 2000; Landys et al., 2006). When facing unpredictable challenges, circulating glucocorticoids increase dramatically, promoting a suite of processes that facilitate responding to and recovering from these challenges (Wingfield et al., 1998; Sapolsky et al., 2000). When sustained, this stress response can trigger an emergency life-history stage in which breeding activities are usually reduced, and energy is redirected toward survival (Wingfield et al., 1998). Thus, glucocorticoids – particularly in the presence of a stressor – have been widely predicted to mediate life history trade-offs between current and future reproduction.

While models of the stress response make clear predictions about redirecting effort, empirical studies have proved equivocal in linking environmental conditions to appropriate responses. One limitation of much empirical work to date is that it has focused on the relationship between glucocorticoids and fitness in a single context (i.e., a single population, year, or environment; Hau et al., 2016; Guindre-Parker, 2018). The factors that shape these relationships are best understood by measuring glucocorticoids across different environmental and life-history contexts (Apfelbeck et al., 2017; Schoenle et al., 2018; Vitousek et al., 2019a). Because the costs and benefits of mounting a robust glucocorticoid stress response are likely to differ across environments and species, populations are predicted to differ in how they regulate glucocorticoids (Schoenle et al., 2018; Vitousek et al., 2019b). For instance, a comparative study in birds found a positive association between latitude and the glucocorticoid response to a standardized retraint stressor (acute challenge that activates the HPA axis resulting in stress-induced glucocorticoids increase) during breeding (Bókony et al., 2009). However, willow warblers (*Phylloscopus trochilus*) breeding in northern Sweden—where the breeding season is shorter and environmental conditions less predictable—show lower stress-induced glucocorticoid levels than those breeding in southern Sweden, where conditions are more predictable (Silverin et al., 1997). Breuner and colleagues (2003) compared males from three populations of white-crowned sparrows (*Zonotrichia leucophrys*) breeding at different latitudes from California to Alaska. They found that males had similar stress-induced glucocorticoid levels, but differed in their corticosteroid-binding globulin and intracellular receptor affinity, suggesting that HPA axis regulation varied across the populations. Thus, while the stress response appears to vary with latitude, additional research is needed to determine which of the many potential selective pressures that covary with latitude are driving this variation.

Two factors thought to be particularly important in shaping the costs and benefits of stress responsiveness are the frequency with which organisms face major unpredictable changes in their environment (the “environmental unpredictability” hypothesis), and how valuable each breeding attempt is to lifetime reproductive effort (the “brood value” or “reproductive value” hypothesis). The “environmental unpredictability” hypothesis predicts that species or populations in less predictable environments will have higher stress-induced glucocorticoid levels to facilitate responding effectively to challenges (Wingfield, 2013; Vitousek et al., 2019a). The reproductive value hypothesis predicts that the stress response differs based on the proportion of lifetime reproductive effort represented by a single breeding attempt. According to this hypothesis, because high glucocorticoid levels can be deleterious to reproduction by diverting energy away from breeding activities, organisms engaging in more valuable reproductive attempts should have lower stress-induced glucocorticoids to avoid jeopardizing the current breeding attempt (Bókony et al., 2009). The relative value of a single reproductive event depends on both the length of the breeding season, which influences the number of potential reproductive attempts an individual can engage in per season, and its lifespan. A shorter breeding season results in less opportunity to engage in multiple successive reproductive attempts or to restart a breeding attempt in case of failure of the initial one, which increases the relative value of each reproductive attempt (Breuner et al., 2003; Wingfield and Sapolsky, 2003; Hau et al., 2016). Therefore, the reproductive value hypothesis predicts that populations living in environments with shorter breeding seasons will show lower stress-induced glucocorticoid levels during breeding (Bókony et al., 2009; Breuner, 2011). This desensitization of the HPA axis to challenges facilitates not interrupting valuable breeding attempts (Wingfield and Sapolsky, 2003; Breuner, 2011).

Previous comparative studies investigating the role of environmental predictability in shaping glucocorticoid concentrations have predominantly used latitude as a proxy for environmental predictability or harshness (e.g., Bókony et al., 2009; Jessop et al., 2013). Variability in environmental conditions, especially temperature, typically increases with increasing latitude and elevation (Breuner 2011); however, these patterns are not always consistent. Furthermore, as many factors (including the duration of the breeding season) also commonly covary with latitude, analyses that use latitude alone could conflate the selective pressures imposed by living in an unpredictable environment with other selective pressures. We are aware of only one previous study that has directly quantified and tested the role of environmental variation in predicting variation in stress-induced glucocorticoids. A phylogenetic comparative analysis of the relative support for these and other factors in shaping glucocorticoid variation across vertebrates found that reproductive value better predicted large-scale variation in stress-induced glucocorticoids than environmental variability (Vitousek et al., 2019a). However, the relative roles of different selective pressures in shaping glucocorticoid variation are expected to vary across species. Disentangling the relative roles of reproductive value and environmental unpredictability in shaping glucocorticoid regulation will require comparing populations of the same species inhabiting different environments.

Central to the hormonal mediation of life history is an understanding of the potential cost of the stress response. The costs of the stress response likely depend not only on maximum glucocorticoid levels but also on the duration of exposure to high levels, which is influenced by the strength of negative feedback. Negative feedback is triggered after activation of the HPA axis and is coordinated by glucocorticoids binding to receptors in the brain inducing a decrease in circulating glucocorticoids (de Kloet et al., 1998; Breuner and Orchinik, 2001; Romero, 2004). Despite increasing evidence that differences in the strength of negative feedback affect aspects of health and performance (Sapolsky, 1983; Dallman et al., 1992; Weaver et al., 2004), its functional effects have been largely neglected in free-living organisms (but see (Romero and Wikelski, 2010; Zimmer et al., 2019). We are not aware of any previous studies that have assessed how negative feedback varies across environments. We hypothesized that differences in negative feedback efficacy could either mitigate or magnify the costs of a stress response. Strong negative feedback could serve to mitigate the costs of mounting a robust stress response by inducing a fast decrease in circulating glucocorticoids. This could be particularly important when a strong stress response is required to cope effectively with frequent unpredictable short-term challenges. In this case, strong negative feedback may make it possible to avoid the negative effects of sustained glucocorticoid elevation and hence recover quickly and resume critical activities such as breeding. A recent study within a single population of tree swallows (*Tachycineta bicolor*) supported the mitigating hypothesis: incubating females that exhibited both a robust stress response and strong negative feedback were less likely to abandon reproductive attempts when facing stressors (Zimmer et al., 2019). Alternatively, coupling an elevated stress response with weak negative feedback could serve to magnify the effects of the stress response. This could be adaptive if mounting a longer stress response facilitates avoiding or alleviating severe challenges; for example, if a more robust response enhances sensitivity to environmental cues (Wingfield, 2013; Vitousek et al., 2019b). Accordingly, individuals breeding in highly variable environments are predicted to show elevated stress repsonses followed by strong negative feedback.

Here, we compared the support for two sets of predictions about how HPA axis regulation differs across populations and environments in breeding tree swallows. First, we asked whether variation in the glucocorticoid stress response (peak stress-induced glucocorticoids) across populations is better predicted by the reproductive value hypothesis or the environmental unpredictability hypothesis. Second, we assessed whether negative feedback varies across populations, and if so, whether the patterns suggested a magnifying or mitigating effect on the stress response. The tree swallow, a common passerine bird that breeds across much of North America, is an ideal species in which to test these predictions as it breeds along an expansive latitudinal as well as elevational gradient within the temperate zone. As such, different populations face different amounts of time available for reproduction as well as differing levels of environmental unpredictability. We compared populations breeding in Tennessee, New York, Wyoming and Alaska. We assessed females’ HPA axis activity by measuring baseline glucocorticoids, stress responses, and negative feedback during two life history stages: incubation and nestling rearing. In order to test whether between-population differences in the stress response are better predicted by differences in the time available for reproduction or environmental unpredictability, we characterized both parameters in each population. We first determined breeding synchrony and the length of the breeding season in each population. Breeding season length is a good proxy for differences in reproductive value across populations breeding at different latitudes (Breuner, 2011). Because increasing reproductive value is generally associated with higher parental investment (Williams, 1966; Ardia, 2005; Ardia and Clotfelter, 2007) and thus potentially with higher reproductive success, we also determined breeding effort and success. To characterize differences in environmental predictability, we calculated an index of unpredictability for different weather variables using historical weather data for each site. We predicted that if variation in the magnitude of the glucocorticoid stress response is predominantly shaped by reproductive value, females breeding at the two sites with a relatively short breeding season (Alaska and Wyoming) would mount a lower acute stress reponse (Bókony et al., 2009; Vitousek et al., 2019a). Conversely, if environmental unpredictability plays a greater role, we expected the opposite pattern: females breeding in the more unpredictable environments of Alaska and Wyoming would have higher stress-induced corticosterone levels. Concerning negative feedback, the mitigation hypothesis predicts strong negative feedback in populations with greater stress responses. The magnifying hypothesis, in contrast, predicts weaker feedback in the face of greater stress responses.

Differences in HPA axis regulation across populations could also be associated with downstream physiological costs. Glucocorticoids affect both redox balance and glucose metabolism (Sapolsky et al., 2000; Costantini et al., 2011). We also tested whether oxdidative stress, another potential physiological mediator of the trade-off between reproduction and survival (Casagrande and Hau, 2018), or glucose levels varied across populations. We predicted that oxidative stress and circulating glucose would be higher in populations breeding in more unpredictable environments and/or with higher reporductive value, and in those with higher baseline glucocorticoid levels.

## Results

### Environmental unpredictability

The unpredictability of temperature during the breeding season was the lowest in Tennessee and the highest in Wyoming and Alaska (Table 1, Fig 1). Temperature unpredictability in New York was intermediate, but closer to Alaska and Wyoming than to Tennessee (Table 1).

**Figure 1:**
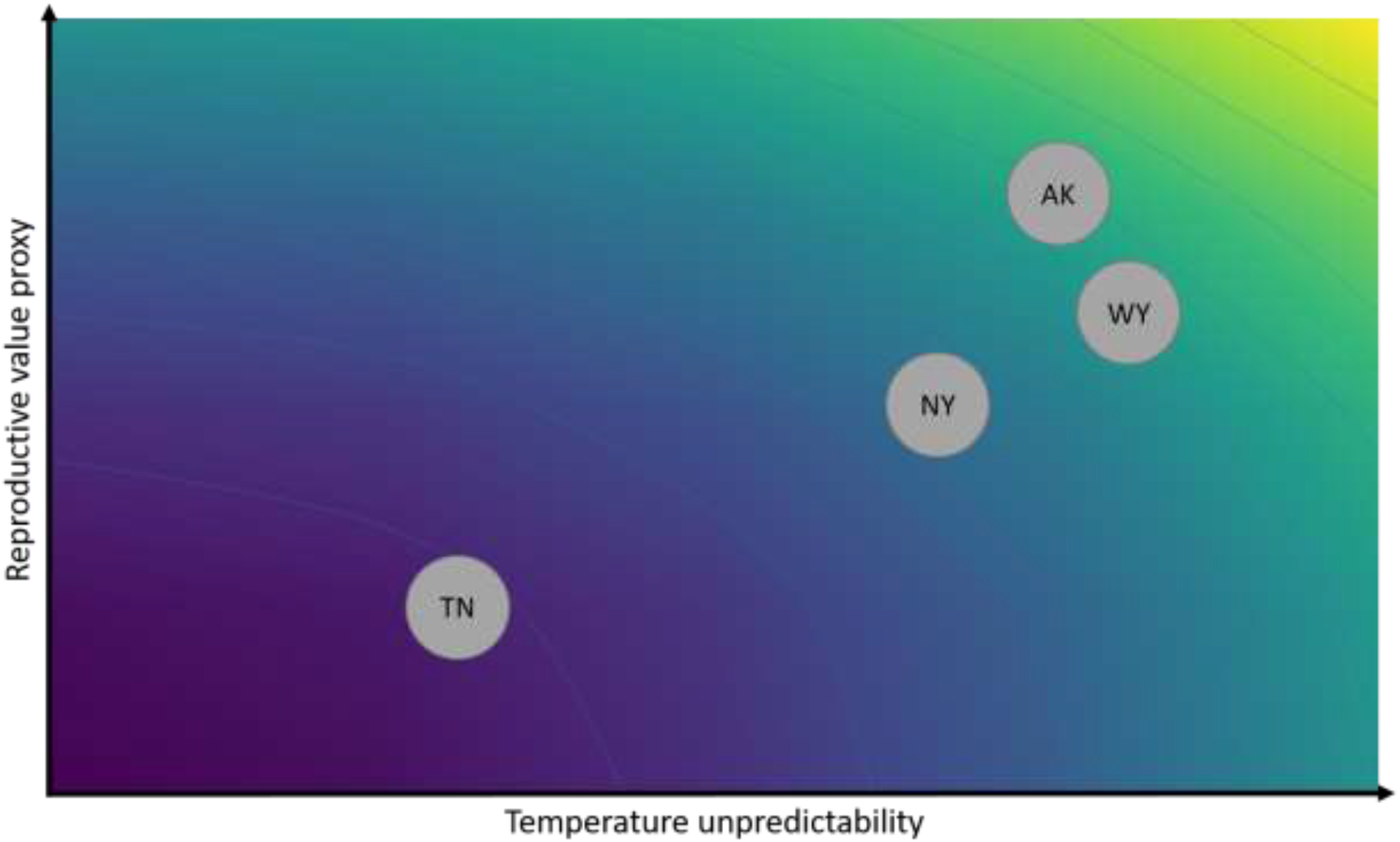
Plot illustrating the difference between the four populations (Tennessee (TN), New York (NY), Wyoming (WY) and Alaska (AK) along a gradient of increasing temperature unpredictability (x axis) and a proxy of reproductive value based on breeding season length (y axis). Warmer color indicates higher environmental unpredictability and higher reproductive value.

**Table 1:** Unpredictability of daily average temperature, daily average active time (between 0600 and 2000) temperature, daily maximum temperature and daily total precipitation at the field sites in Tennessee, New York, Wyoming and Alaska. Higher SD_res_ value indicates greater unpredictability.

|  | Tennessee | New York | Wyoming | Alaska |
| --- | --- | --- | --- | --- |
| $SD_{res}$ average daily temperature | 0.18 | 0.33 | 0.41 | 0.38 |
| $SD_{res}$ average daytime temperature | 0.21 | 0.42 | 0.53 | 0.50 |
| $SD_{res}$ maximum daily temperature | 0.21 | 0.38 | 0.51 | 0.47 |

### Reproductive value

The synchrony with which females laid first clutches differed between populations (χ^2^_3,250_ = 89.51, p < 0.0001) with females in TN (17.7 ± 1 days) showing a more extended clutch initiation period than females in the three other populations (NY: 8.9 ± 1.1 days, WY: 9.2 ± 0.7 days, AK: 8.8 ± 0.6 days; z ≥ 6.68, p < 0.0001). Total length of the breeding season also differed between populations (χ^2^_3_ = 8.49, p = 0.037) with a longer breeding season in TN (99 days) and the shortest in AK (66 days). Breeding season lengths in NY (74 days) and WY (70 days) were intermediate, but more similar to AK than TN. Overall, these patterns suggest that reproductive value was the lowest in TN and relatively similar across the other three populations (Fig 1).

### Corticosterone regulation

Females’ corticosterone phenotypes differed between populations. This difference was influenced by the life history substage during which the female was captured and corticosterone sample type (baseline, stress-induced, post-dexamthasone (post-dex); population x life history substage x sample: F_12,1263_ = 3.92, p < 0.0001; Fig 2).

**Figure 2:**
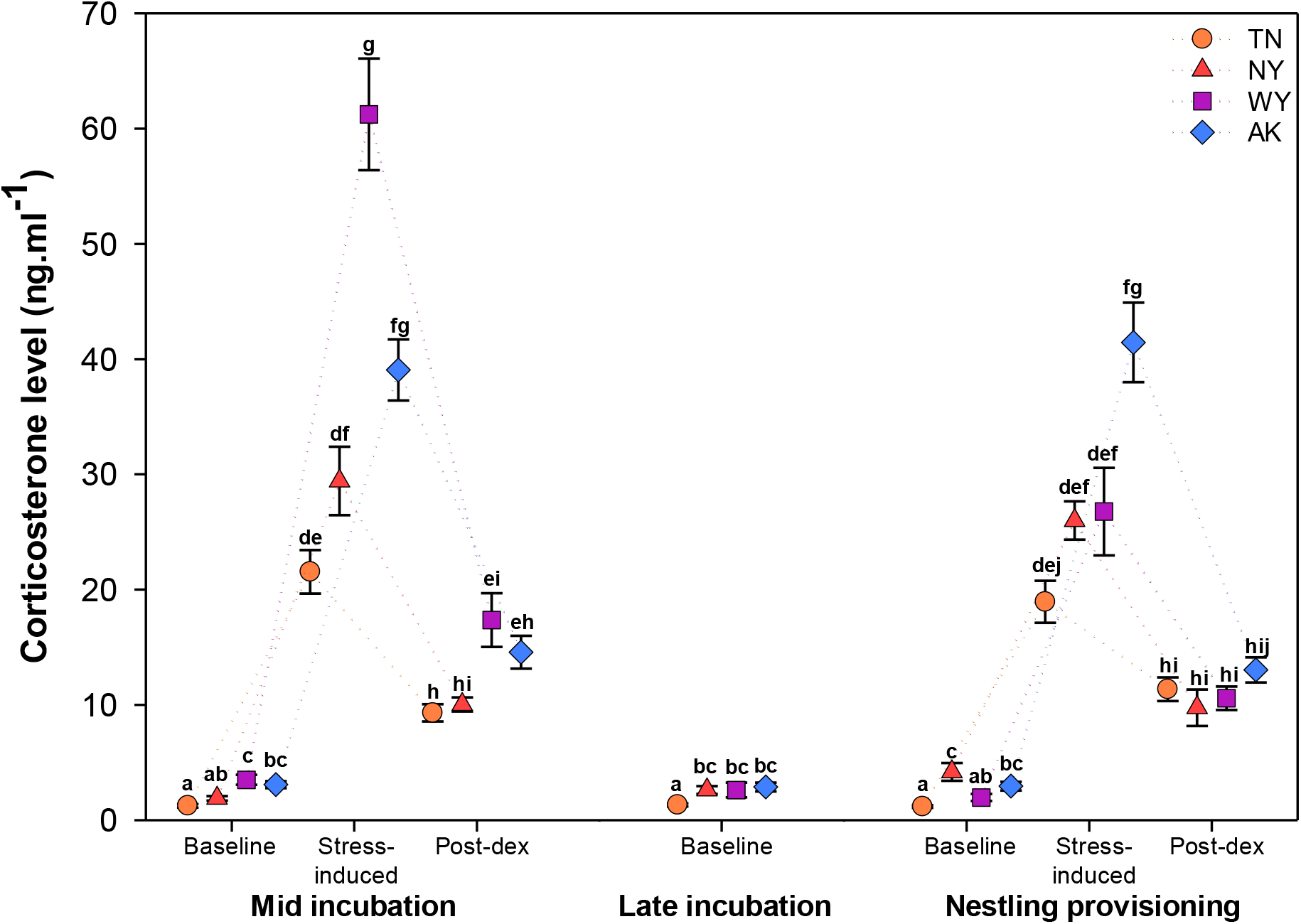
Corticosterone levels (mean ± SE) of females when captured at three points during the reproductive cycle (mid incubation: day 6-7 of incubation, late incubation: day 12-13 of incubation, and nestling provisioning: day 6 post-hatch) in Tennessee (TN), New York (NY), Wyoming (WY), and Alaska (AK). At all captures baseline corticosterone levels were measured. During the first and third captures, stress-induced corticosterone after 30 minutes of restraint and 30 minutes after injection of dexamethasone (post-dex) were also measured. Different letters indicate significant differences.

Across sampling periods, baseline corticosterone levels tended to be lower in Tennessee than in the other three populations (Fig. 2, see supplementary material for statistical analyses).

Circulating stress-induced corticosterone levels generally increased with temperature unpredictability (Fig 2). The exception to this pattern was in WY during mid incubation, when stress-induced corticosterone levels were significantly higher than all other populations except for AK (see supplementary material for statistical analyses). Corticosterone stress responses (the difference between stress-induced and baseline corticosterone) showed a similar pattern. Overall, stress responses differed between populations and life history substages (population x life history substage: F_3,220.1_ = 9.45, p < 0.0001; Fig 3a). During mid-incubation, stress responses were highest in WY (t ≥ 5.08, p < 0.0001; Fig 3a) and intermediate in AK (t ≥ 2.71, p ≤ 0.044; Fig 2a). During nestling provisioning, females in AK had a stronger stress response than females in all other populations (t ≥ 3.01, p ≤ 0.048; Fig 3a).

**Figure 3:**
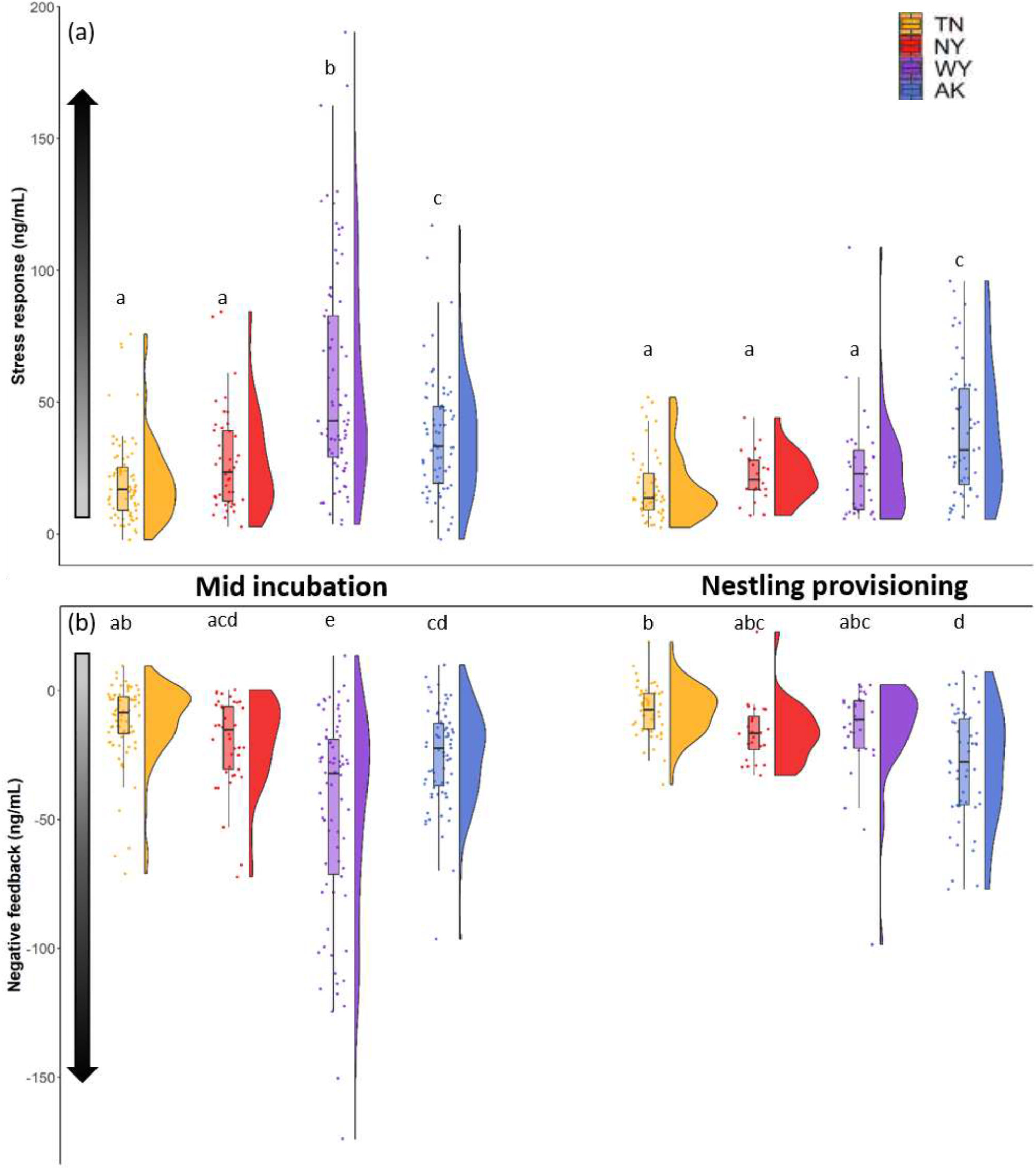
Females’ glucocorticoid stress responses (a) and the strength of negative feedback (b) in Tennessee (orange), New York (red), Wyoming (purple) and Alaska (blue) during mid incubation and nestling provisioning. Points overlaying the boxplot show raw data and the half-split violin shows the probability density function. Shaded arrows along the y-axis denote increasing strength of the stress response and negative feedback with increasing darkness. Different letters indicate significant differences.

Circulating corticosterone levels after dexamethasone injection did not differ between the four populations (t ≤ 1.73, p ≥ 0.99; Fig 2). However, negative feedback, i.e., the decrease in corticosterone following dexamethasone injections, differed between populations and life history substages (population x life history substage: F_3,236.3_ = 8.30, p < 0.0001; Fig 3b). During mid-incubation, negative feedback was stronger in WY than in all other populations (t ≥ 5.36, p < 0.0001; Fig 3b). In the three other populations, negative feedback was highest in AK, intermediate in NY, and weakest in TN (pairwise comparisons: AK vs. TN: t = 3.60, p = 0.016, AK vs. NY: t = 0.69, p = 0.99, TN vs. NY: t = 1.55, p = 0.77; Fig 3b). During the nestling provisioning period, females in AK had stronger negative feedback than females in all other populations (t ≥ 3.16, p ≤ 0.04; Fig 3b); negative feedback did not differ among the other populations (t ≤ 1.24, p ≥ 0.91; Fig 3b). Overall, negative feedback was stronger in populations that mounted a strong stress response.

Within populations, the magnitude of the stress response did not change between substages of the reproductive period (t ≤ 1.49, p ≥ 0.81; Fig 3a), except in WY where it decreased between incubation and nestling provisioning (t = 7.09, p < 0.0001; Fig 3a). Negative feedback efficacy also did not change between life history substages within populations (t ≤ 1.28, p ≥ 0.90; Fig 3b), except in WY where it decreased between mid-incubation and nestling provisioning (t = 6.08, p < 0.0001; Fig 3b). See supplementary material for detailed information about within population corticosterone changes across the breeding season.

Correlations between the different measures of the HPA axis differed between populations and life history substages. Overall, the different aspects of the HPA axis are not correlated or weakly to moderately correlated(Table 2). However, the strength of the stress response and the efficacy of negative feedback are positively correlated in all populations at both life history substages (Table 2).

**Table 2:** Correlations between baseline, stress-induced and post-dexamethasone (post-dex) glucorticoid levels and between the strength of the stress response and the efficacy of negative feedback within each population. Data are R^2^, p-value. Bold indicates significant correlations.

|  |  | Mid incubation |  |  | Nestling provisioning |  |  |
| --- | --- | --- | --- | --- | --- | --- | --- |
|  |  | stress-induced<br>(R<sup>2</sup>, p-value) | post-dex<br>(R<sup>2</sup>, p-value) | negative<br>feedback<br>(R<sup>2</sup>, p-value) | stress-induced<br>(R<sup>2</sup>, p-value) | post-dex<br>(R<sup>2</sup>, p-value) | negative<br>feedback<br>(R<sup>2</sup>, p-value) |
| Tennessee | baseline | 0.001, 0.94 | 0.005, 0.54 |  | **0.09, 0.03** | **0.18, 0.002** |  |
|  | stress-induced |  | **0.08, 0.02** |  |  | **0.27, 0.0001** |  |
|  | stress response |  |  | **0.84, <0.0001** |  |  | **0.70, <0.0001** |
| New York | baseline | 0.02, 0.43 | 0.02, 0.41 |  | 0.001, 0.89 | 0.03, 0.43 |  |
|  | stress-induced |  | **0.15, 0.01** |  |  | 0.01, 0.62 |  |
|  | stress response |  |  | **0.96, <0.0001** |  |  | **0.50, 0.0008** |
| Wyoming | baseline | **0.07, 0.02** | 0.005, 0.55 |  | **0.11, 0.07** | 0.01, 0.60 |  |
|  | stress-induced |  | 0.001, 0.91 |  |  | 0.08, 0.12 |  |
|  | stress response |  |  | **0.91, <0.0001** |  |  | **0.92, <0.0001** |
| Alaska | baseline | 0.001, 0.78 | 0.003, 0.66 |  | 0.01, 0.43 | 0.007, 0.54 |  |
|  | stress-induced |  | 0.03, 0.15 |  |  | **0.25, 0.0001** |  |
|  | stress response |  |  | **0.69, <0.0001** |  |  | **0.90, <0.0001** |

### Reproductive effort and success

There were no differences in clutch size or brood size at hatching between populations (χ^2^ ≤ 5.90, p ≥ 0.12, see supplementary material). Because of an extended period of cold, wet weather that occurred during the incubation stages of most females, hatching success (χ^2^_3,250_ = 29.24, p < 0.0001) was lower in WY (50.7 %, 38 of 75 nests) than in the other three populations (TN: 82.2 %, 60 of 73 nets; NY: 79.5 %, 31 of 39 nests; AK: 84.4 %, 54 of 64 nests; z ≥ 3.58, p ≤ 0.002). Provisioning effort differed across populations (F_3,155.4_ = 6.00, p = 0.0007). Females in AK and WY provisioned at higher rates than those in NY and TN (Fig 4).

**Figure 4:**
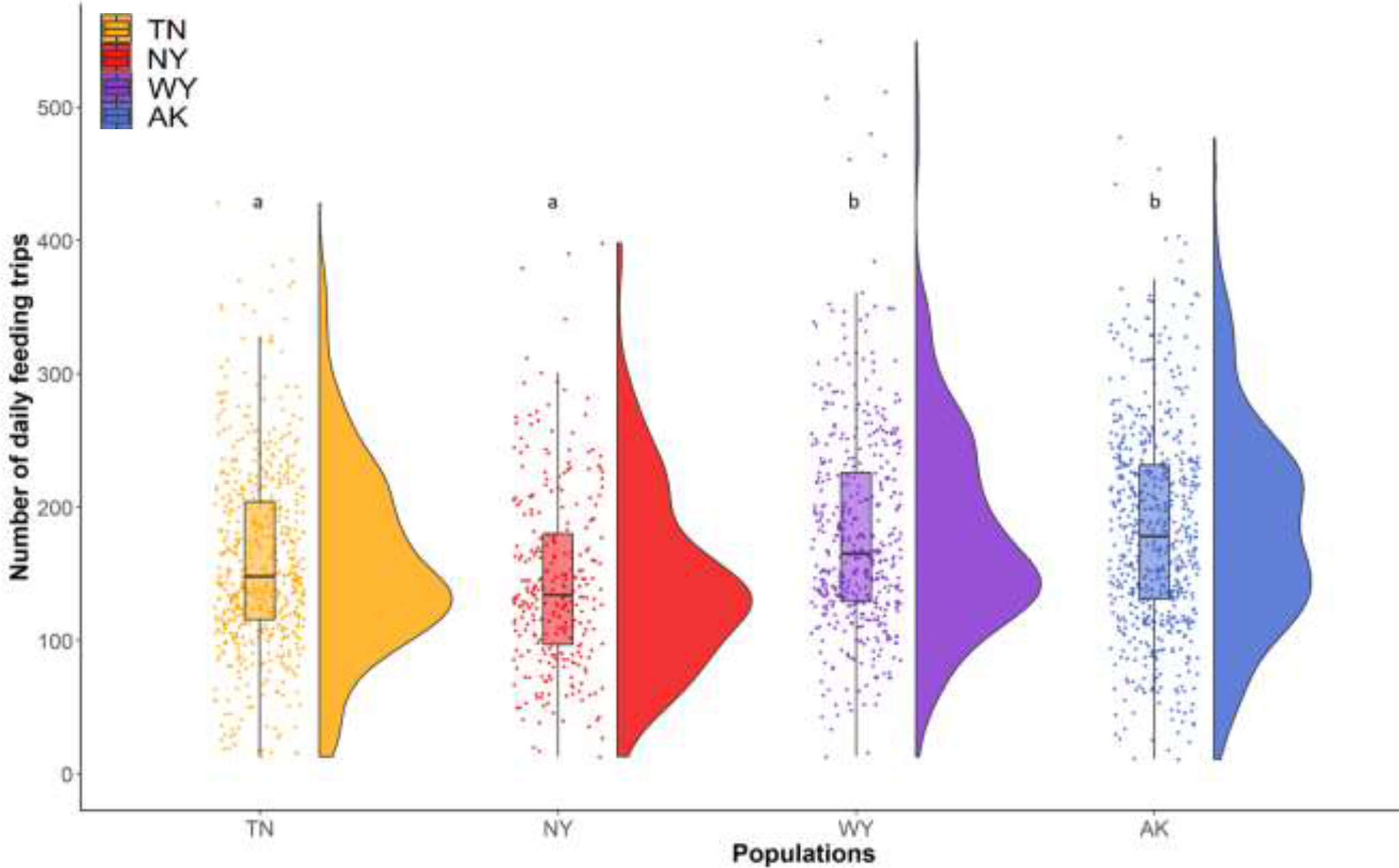
Number of daily feeding trips for females in Tennessee (orange), New York (red), Wyoming (purple), and Alaska (blue), calculated from the total number of raw visits recorded (329,047). Points overlaying the boxplot show raw data and the half-split violin shows the probability density function. Different letters indicate significant differences between populations.

Nestling body mass differed across populations, mirroring the observed differences in female provisioning rates (F_3,128.5_ = 4.02, p = 0.009). On day 12 post-hatching, nestlings were heaviest in AK (21.2 ± 0.2 g) and WY (20.6 ± 0.2 g), and lightest in NY (18.7 ± 0.5 g) and TN (19.6 ± 0.2 g). Among nests in which one or more eggs hatched, the number of nests in which one or more nestlings fledged also differed among populations (χ^2^_3,250_ = 22.49, p < 0.0001); fledging success was lower in NY (35.5 %, 11 of 31 nests) than in the other populations (TN: 76.7 %, 46 of 60 nests, AK: 80.4 %, 41 of 51 nests, WY: 80.0 %, 28 of 35 nests; t ≥ 3.30, p ≤ 0.005).

## Discussion

In the current context of global changes, understanding how organisms have evolved in response to environmental challenges is critical. Our results demonstrate that glucocorticoid regulation across different environmental and life-history contexts may underlie successful adaptation and population persistence. First, we showed that female tree swallows breeding in environments with higher temperature unpredictability secrete more glucocorticoids in response to acute challenges during breeding. This pattern was strong even though at these sites birds also experienced shorter breeding seasons and would normally be predicted to skew investment towards offspring. Thus, if elevated glucocorticoids triggered reproductive failure or delay, birds at those sites would have less time to complete reproduction and re-nest if necessary. This seemingly contrary result reveals environmental unpredictability may be a stronger force than the value of the current breeding attempt in determining stress-induced glucocorticoid levels in female tree swallows. Second, we found that populations with higher stress responses also have stronger negative feedback, which likely functions to limit potential damage caused by high glucocorticoid levels. The combination of high stress-induced glucocorticoid levels followed by the induction of strong negative feedback may allow females to cope effectively with frequent unpredictable challenges, and to recover faster in order to continue breeding activities. This phenotype may be most strongly favored in environments with both high unpredictability and greater time constraints on reproduction.

### Life history and environmental unpredictability

Across the four study populations we found differences in key selective pressures. Birds breeding in Alaska and in the mountains of Wyoming faced more unpredictable temperatures than birds in Tennessee; New York conditions were intermediate. The time constraints on reproduction differed similarly across populations, resulting in higher apparent reproductive value in Alaska and Wyoming and lowest apparent reproductive value in Tennessee. As predicted by life history theory (and consistent with previous comparative work in tree swallows: Ardia, 2006; Ardia, 2007; Rose and Lyon, 2013; Akçay et al., 2016), birds in Alaska and Wyoming showed higher parental investment (measured as offspring feeding rate) and reared larger offspring than those in New York and Tennessee.

### Glucocorticoid regulation

At our two sites with stronger time constraints on reproduction and high temperature unpredictability (AK and WY) we found that the magnitude of the glucocorticoid stress response was greatest. This result provides the first empirical support for the idea that environmental unpredictability could be a stronger force shaping the peak glucocorticoid response to challenges than brood value, which should select for suppressed stress responses in the face of low residual reproductive value. In contrast with this finding, a recent large scale phylogenetic comparative analysis found greater support for reproductive value shaping glucocorticoid stress responses across vertebrates than environmental variation (Vitousek et al. 2019b). Although other studies have not directly compared these effects, a number of analyses have supported reproductive value as a driver of variation in the strength of glucocorticoid stress responses within and across species (Silverin et al., 1997; Bókony et al., 2009; Breuner, 2011; Hau et al., 2016, but see Breuner et al., 2003; Krause et al., 2016). However, previous studies have also found that latitude (which is often used as a proxy for environmental variation or harshness: e.g., Bókony et al., 2009; Jessop et al., 2013) covaries positively with stress-induced glucocorticoids. We predict that differences in the relative importance of environmental unpredictability and reproductive value in shaping stress responsiveness is likely to vary across species. Specifically, we predict that the stress responsiveness of income breeders will be more strongly influenced by environmental unpredictability than that of capital breeders. Similarly, we predict that stress responsiveness will be particularly closely tied to environmental unpredictability in species that rely on critical food sources impacted by short-term environmental fluctuations (e.g., aerial insectivores).

A robust stress response could allow appropriate responses to challenges, but might also result in associated reproductive costs (Wingfield and Sapolsky, 2003). However, we found no evidence that maintaining a robust stress response had negative effects on reproduction. Reproductive success did not vary consistently across populations; instead, the differences that we saw in hatching success (lowest in WY) and fledging success (lowest in NY) mirrored temporary periods of inclement weather in those populations. Consistent with life history theory, females in Alaska and Wyoming showed a higher investment in nestlings. Nevertheless, it is possible that the costs of maintaining an elevated stress response would manifest under more prolonged stressful conditions, or that it imposes longer-term costs (e.g., accelerated telomere shortening or senescence).

We found support for a mitigating effect of negative feedback on stress responsiveness: female tree swallows breeding in Alaska and Wyoming, where both time constraints on reproduction and temperature unpredictability were the highest, showed both a high stress response and strong negative feedback. Stronger negative feedback allows for a faster decrease in circulating glucocorticoids, potentially reducing the costs associated with sustained glucocorticoid elevation. Thus, strong negative feedback should be particularly beneficial for individuals with a strong stress response (Zimmer et al., 2019). We recently demonstrated that variation in negative feedback efficacy is related to the speed of recovery of the HPA axis from repeated transient stressors, stress resilience, and reproductive success in tree swallows (Taff et al., 2018; Vitousek et al., 2019b; Zimmer et al., 2019). These results indicate that regulation of negative feedback is crucial for coping with perturbances, particularly in more challenging environments.

Taken together, our results suggest that when environmental conditions become more variable female tree swallows couple high magnitude stress responses and strong negative feedback, instead of decreasing the hormonal response to challenges. Through strong negative feedback individuals could limit the reproductive costs of glucocorticoid exposure, and resume reproductive activities as soon as the stressor has passed. Thus, this combination of glucocorticoid regulatory traits may enable individuals living in harsher environments to balance the challenges of high environmental unpredictability and a brief reproductive period in order to maximize survival and reproductive success.

Our findings are also consistent with the idea that interactions among glucocorticoid regulatory elements may be important for appropriately responding to challenges, and for fitness (Vitousek et al., 2018b; Vitousek et al., 2019b; Zimmer et al., 2019). In all four populations the strength of the stress response and the efficacy of negative feedback positively covary; however, there were weakly positive or no phenotypic correlations between stress-induced and post-dexamethasone corticosterone levels. Covariation in components of glucocorticoid regulation could result from similar regulatory pathways, as negative feedback is regulated by glucocorticoids binding to glucocorticoid receptors (de Kloet et al., 1998; Romero, 2004). As such, higher glucocorticoid levels could activate more receptors, inducing faster negative feedback (Breuner and Orchinik, 2001; Romero, 2004). However, it has been suggested that different components of the HPA axis are modulated independently (e.g., Romero, 2004). Therefore, the phenotypic correlations seen here could also result from selection favoring combinations of these traits. Determining whether stress-induced and post-dexamethasone corticosterone levels are genetically correlated and whether glucocorticoid profiles covary with receptor expression could help to illuminate the flexibility and physiological underpinnings of these traits and their potential to respond to selection.

Baseline glucocorticoid levels also differed across populations. However, as both the reproductive value hypothesis and the environmental unpredictability hypothesis predict higher baseline glucocorticoid levels in Alaska and Wyoming, where the season is short and weather unpredictable, these patterns do not allow us to differentiate between the role of these forces in shaping glucocorticoid evolution. Elevated baseline glucocorticoid levels have been shown to support energetically demanding activities, and are associated with more challenging conditions and increased investment in reproduction within and across species (Hau et al., 2010; Bonier et al., 2011; Jessop et al., 2013; Apfelbeck et al., 2017; Vitousek et al., 2019a). The particularly high baseline (and stress-induced) corticosterone levels in Wyoming during incubation may also have reflected a temporary upregulation in HPA axis activation because of a period of unusually cold and wet weather that occurred during this time. In contrast, in Tennessee, in which tree swallows experience a long breeding season and more predicatable conditions baseline glucocorticoid levels were consistently low throughout the reproductive period.

Contrary to our predictions that elevated stress levels would incur physiological costs, we did not detect differences between populations in measures of oxidative damage, antioxidant capacity, or glucose metabolism. Investment in reproduction is not always associated with higher oxidative stress (Garratt et al., 2011; Speakman and Garratt, 2014). However, within individuals both oxidative stress and glucose have been shown to increase with increasing glucocorticoids levels or workload (Sapolsky et al., 2000; Costantini et al., 2011; Metcalfe and Monaghan, 2013). The lack of these relationships at the population level could reflect different trade-offs operating across populations and within individuals.

### Conclusion

These findings provide support for the hypotheses that environmental unpredictability may be a critical factor in shaping glucocorticoid stress responses, and that selection favoring strong negative feedback in more stress responsive individuals could serve as a mechanism to mitigate the costs of mounting a strong stress response. Our results are also in accordance with the hypothesis that negative feedback and the dynamic regulation of glucocorticoids are important for coping with challenging conditions (Romero and Wikelski, 2010; Taff and Vitousek, 2016; Vitousek et al., 2019b). In the current context of global changes, intraspecific differences in the response to stressors may be particularly important for survival or for the ability to adapt to new conditions (Angelier and Wingfield, 2013; Harding et al., 2019). Our results suggest that this phenotype (elevated stress response and strong negative feedback) has been selected for in unpredictable environments and might therefore be expected to become increasingly common over time, assuming genetic variation exists. However, as climate change affects both the length of the breeding season (Dunn and Winkler, 2010) and environmental predictability (Thornton et al., 2014), populations may face rapidly changing regimes of selection on glucocorticoid regulation outside the bounds of evolutionary history. Confirming that selection is occurring in these populations will require testing whether among individual differences in glucocorticoid phenotype affect fitness. Ultimately, determining the evolutionary causes and consequences of differences in glucocorticoid levels within and among populations will help to reveal how selection drives HPA axis regulation and whether the history of selection on hormonal regulation influences the ability to cope with unpredictable or changing environments.

## Methods

### Populations

Field data were collected from 2016 to 2018 in four different populations of tree swallows breeding in nest-boxes. Populations were located in Chattanooga, Tennessee (TN) (35.1°N, 85.2°W, 206m elevation), Ithaca, New York (NY) (42.5°N, 76.5°W, 340m elevation), Burgess Junction, Wyoming (WY) (44.5°N, 107.3°W, 2451m elevation) and in McCarthy, Alaska (AK) (61.4°N, 143.3°W, 445m elevation). Tree swallows are widely distributed across North America and breed in a variety of environments. These populations were chosen to allow for comparisons among populations breeding in environments with different degrees of environmental predictability and reproductive value. Populations at higher latitude (Alaska) or elevation (Wyoming) are expected to experience cooler and more unpredictable weather conditions and a shorter breeding season starting later in the year (late May). In Tennessee, near the Southern edge of the breeding distribution, tree swallows are expected to experience warmer and more predictable weather conditions and a long breeding season with the first egg usually laid earlier in the season (early to mid-April). In New York, environmental conditions and breeding season length are expected to be intermediate, with the first egg usually laid in early May.

### General field methods and stress manipulation

In the four populations, nests were monitored every 1-2 days throughout the breeding season from the initiation of activity at each site to fledging, except for the last week of nestlings’ development (to avoid inducing premature fledging). For every active nest, we recorded clutch initiation date and completion dates, clutch size, hatch date, brood size, and the number of nestlings fledged. Nestling fates were determined by checking boxes 22-24 days after hatching. We installed radio frequency identification (RFID) units on each box on the fourth day of incubation (see below). Birds were captured at their nest boxes by hand or using a manually activated trap. All birds were captured and sampled on specific days of life history substages, and during a set time of day, to reduce the variation in circulating glucocorticoid hormones resulting from circadian rhythms. Adult females were captured between 0700 and 1000h in NY, TN and WY and between or 0600 and 0900h in AK to compensate for the earlier start of activity due to the increased day length compared to the other populations.

Females were initially captured 6 or 7 days after clutch completion (capture number 1). At this capture, we took a first blood sample within 3 min of initial disturbance to measure baseline circulating corticosterone levels. A second blood sample was taken after 30 min of restraint in a cloth bag to measure stress-induced corticosterone levels. Immediately after this sample was taken, females were injected with dexamethasone (dex) (0.5 μl.g^-1^, Dexamethasone Sodium Phosphate, Mylan Institutional LLC), a synthetic glucocorticoid that binds to receptors within the HPA axis, in order to induce negative feedback (Zimmer et al., 2019). A final blood sample was taken 30 min after dex injection to measure the degree of down-regulation in circulating corticosterone (a measure of negative feedback). Between samples, we weighed the females, and measured the length of their skull from the back of the head to the bill tip (head-bill) and flattened wing length. Non-banded individuals received USGS leg bands and a celluloid color band with attached passive integrated transponder (PIT) tag encoding a 10-digit hexadecimal string (Cyntag, Cynthiana, KY). Female age was determined based on plumage coloration and characterized as second year (SY) or after second year (ASY) (Hussell, 1983).

As part of a separate study, at their first capture, adult females were allocated to one of the experimental groups: control, feather restraint (in which three primaries were reversibly attached to alter flight ability and thereby increasing the cost of foraging), or predator exposure (see Zimmer et al., 2019 for details on both experimental treatments). Treatments started after the first capture and lasted for 5-6 days.

Females were then recaptured 5-6 days later (on incubation day 12 or 13; capture 2), at the end of the experimental treatments (see above). At this capture, we only took a baseline blood sample and weighed the bird before release. Finally, we recaptured females again 6-8 days after eggs hatched (capture 3). We followed the same procedure as in capture 1, taking a baseline, restraint stress-induced, and post-dex blood samples, and again weighed each female.

Twelve days after eggs hatched, each nestling received an USGS leg band, was weighed and had head-bill and flat wing length measured.

All blood samples were collected from the alar vein, in heparinized microhematocrit capillary tubes. Glucose levels in baseline and stress-induced blood samples were determined in the field (see supplementary). Blood samples were then transferred to microcentrifuge tubes, and kept on ice until centrifugation (within 4h). After separation, the plasma was stored at −20°C in the field and then at −80 °C in the lab until analysis. All methods were approved by Cornell IACUC and conducted with appropriate state and federal permits.

### Provisioning behavior

Number of feeding trips for females from nestling ages 1-18 was recorded using radio-frequency identification (RFID) devices (Cellular Tracking Technologies; Rio Grande, NJ, USA) (Bridge and Bonter, 2011). RFID units were installed on each active box on day 4 of incubation. Antennae were fastened around each entrance hole so that birds had to pass directly through an antenna to enter or exit the box. We programmed our RFID units to sample for PIT tags every second between 0500 and 2200h each day as tree swallows are not very active at night. Poll time was set at 500, and cycle time at 1000. The delay time (minimum period of time between successive tag recordings) was set to 1s. RFID boards were powered by 12V 5Ah (PS-1250, PowerSonic, San Diego, CA) batteries that were replaced every five days. At the first capture, each bird was fitted with a PIT tag attached to a color band. Each PIT tag encoded a unique 10-digit hexadecimal string that was recorded, along with a time stamp, when birds passed through or perched on the antenna (see Vitousek et al., 2018a for more details). From the raw RFID records, we determined the number of daily feeding trips for each female through 18 days of age for the brood using an algorithm validated in the New York population (Vitousek et al., 2018a).

### Corticosterone assay

Steroids were extracted from plasma samples using a triple ethyl acetate extraction and then corticosterone levels were determined using an enzyme immunoassay kit (DetectX Corticosterone, Arbor Assays: K014-H5) previously validated for tree swallows (Taff et al., 2019). Samples were run in duplicate and all samples from an individual were run on the same plate. In total we ran 47 assays with an average extraction efficiency of 92.8 % and a detection limit of 0.47 ng.ml^-1^. The intra-assay variation was 8.88 % and the inter-assay variation was 11.1 %.

### Data analysis

To characterize the degree of environmental unpredictability at the different field sites we calculated the unpredictability of temperature variables. We obtained historical weather data for each site over as long a yearly range as possible. For New York we obtained data from the North East Climate Center (http://www.nrcc.cornell.edu/) for the Game Farm road weather station (from 1983, located about 7km from field sites) and from the Western Regional Climate Center (https://wrcc.dri.edu/) for the Prentice Cooper State Forest station in Tennessee (from 2003, located about 16 km from field sites), the May Creek station in Alaska (from 1990 located about 19 km from field sites) and the Burgess station in Wyoming (from 1992, located about 5 km from field sites). From these data, we extracted the average daily temperature, the daytime average daily temperature (between 0600 and 2200) which is the period when the swallows are the most active and the daily maximum temperature, which is known to affect flying insects’ activity and therefore food availability (Winkler et al., 2013).

We quantified the unpredictability of these temperatures variables during the breeding season: from April to June in Tennessee and New York and from May to July in Alaska and Wyoming. We calculated unpredictability using a general additive model (GAM) following the methods described in Franch-Gras et al. (2017). For each site, these variables were divided by their mean to normalize them before analysis (Franch-Gras et al., 2017). This model considers the dispersion of the data in the time series around a typical curve of the normalized variable. For each weather variable, the typical curve was fitted in a GAM model in relation to the day of the year using the gam function in the mgcv package in R 3.5.3 (R Core Team, 2019). As suggested by Franch-Gras (2017), we fitted the GAMs using cubic splines as smoothing function to not *a priori* constrain the shape of the curve. The standard deviation of the residuals of the fitted model (SD_res_) represents an index of unpredictability for each variable (Franch-Gras et al., 2017). This index gives an overall measure of unpredictability using the historical records and is not intended to indicate variation in weather in the particular years of study at each site.

We compared females’ corticosterone levels by fitting a generalized linear mixed model (GLMM) with a gamma distribution that included population, capture number, sample (baseline, stress-induced and post-dex), female age and their interactions as fixed factors, relative clutch initiation date as a covariate and female identity as random factor. We did not include treatment upon first capture (i.e., feather restraint, predator exposure, or control) as an independent variable in the above analyses as it did not affect HPA axis regulation, i.e., corticosterone levels (treatment x sample: F_4,1202_ = 0.60, p = 0.66; treatment x capture number x sample: F_8,1254_ = 1.51, p = 0.15). We further characterized females’ corticosterone regulation by calculating their glucocorticoid stress response as the difference between stress-induced and baseline corticosterone levels and their negative feedback as the difference between post-dex and stress-induced corticosterone levels. We compared the magnitude of females’ acute stress response and negative feedback strength using GLMMs fit with a normal distribution including population, capture number, female age and their interactions as fixed factors, relative clutch initiation date as a covariate and female identity as a random factor. Within each population, we determined whether corticosterone level at each time point, and stress response and negative feedback were correlated using Pearson correlations.

In order to compare the populations in terms of breeding synchronization, we calculated the relative clutch initiation date of the first clutch as the number of days after the first laying female in the population. Then, we compared relative clutch initiation date between populations using GLMs with population, female age and their interaction as fixed factors. We also calculated the total breeding season length as the number of days between the first clutch initiation and the last day nestlings fledged at each site. We compared populations usimg a GLM fitted with a Poisson distribution. We also compared clutch size, brood size, hatching success and fledging success between populations. Population, female age and their interactions were added as fixed factors, and relative clutch initiation day as a covariate. The model for relative clutch initiation date was fitted with a normal distribution, models for clutch size and brood size with a Poisson distribution and models for hatching success and fledging success with a binomial distribution.

We compared the number of daily feeding trips females made using a generalized linear mixed model GLMM fitted with a Poisson distribution. Population, female age, their interactions and nestling age were added as fixed factors and brood size at each nestling age as a covariate. We also added relative clutch initiation date and brood size as covariates and nest identity as random factor. We used GLMMs to compare nestlings’ body mass with population, female age and their interaction as fixed factors. Female identity was added as a random factor.

GLMs were run using the GENMOD procedure and GLMMs the GLIMMIX procedure in SAS University Edition (SAS Institute Inc., Cary, NC). Post-hoc comparisons were performed using Tukey-Kramer multiple comparison adjustment to obtain corrected p-values. Probability levels <0.05 were considered significant. Data are presented as mean ± SE.

## Acknowledgments

Funding was provided by NSF IOS grant 1457151 to MV. We thank the many field and lab assistants who helped who helped with data collection.

