## Supplementary material for "Environmental unpredictability shapes glucocorticoid regulation across populations of tree swallows"

### Methods

*Glucose and oxidative stress measures*

Glucose levels in baseline and stress-induced blood samples were determined using FreeStyle Freedom Lite portable glucose meters (Abbott Diabetes Care, Alameda, USA), which have been validated for use in the field in birds (Malisch et al., 2018).

The d-ROMs test quantifies the concentration of reactive oxygen metabolites that result from the oxidation of biomolecules. Protocols were run according to the manufacturer's instructions, except that reaction volumes were fractionally reduced to fit a 96-well plate. 5µL of plasma were pipetted into the sample wells, 5 µL of a calibrator solution in reference standard wells and 5 µL of distilled water in blank wells. Then, 200µL of acetate buffer mixed with 2µL of chromogen were added to each well. We did a kinetic reading that recorded absorbance at a wavelength of 505 nm every minute for 10 minutes. Concentrations were calculated based on the kit instructions and converted to mM of H2O2 equivalents. Measures of variation were estimated by running pooled plasma samples repeatedly within and across plates. Intra-plate variation was 5.2 % and inter-plate variation was 5.5 %.

The OXY-adsorbent test assesses the ability of plasma antioxidants to resist oxidation by hypochlorous acid (HOCl), an endogenously produced oxidant. Samples were run to the manufacturer's instructions, with reaction volumes fractionally reduced to fit a 96-well plate. Plasma samples (2µL) were diluted 1:50 with distilled water. 5µL of the diluted plasma solution was incubated for 10 min at room temperature with a 200 µL of HOCl solution. 5µL of calibrator solution was pipetted in reference standard wells and 5µL of water was in blank wells, both of which were then incubated with 200µL of HOCl. 2µL of the supplied chromogen solution was added to each well following incubation, and the absorbance was read immediately at a wavelength of 505nm. Concentrations were calculated following the kit instructions and expressed in mM of HOCl neutralized per mL of sample. Measures of variation were estimated by running pooled plasma samples repeatedly within and across plates. Intra-plate variation was 4.1% and inter-plate variation 10.1%.

*Data analysis*

We compared females' corticosterone levels by fitting a generalized linear mixed model (GLMM) with a gamma distribution that included population, capture number (capture 1 during mid-incubation, capture 2 at the end of incubation and capture at day 6-8 of nestling provisioning), sample (baseline, stress-induced and post-dexamethasone injection (post-dex)), female age, and their interactions as fixed factors, relative clutch initiation date (as the number of days after the first laying female in the population ) as a covariate, and female identity as random factor. Within each population, we determined whether corticosterone level at each point, and stress response and negative feedback were correlated using Pearson correlations.

We compared female body mass between the populations and treatments over the 3 captures using a GLMM with population, treatment, capture number, female age, and their interactions as fixed factors. We specified relative clutch initiation date as covariate and female identity as random factor. The model was fitted with a gamma distribution.

We compared glucose level using a GLMM fitted with a gamma distribution that included population, treatment, capture number, sample (baseline and stress-induced), female age, and their interactions as fixed factors, relative clutch initiation date as covariate, and female identity as random factor. Finally, we compared OXY and dROMS levels using the previous model excluding sample as factor as we only determined baseline levels.

For each nestling, we calculated their scaled body mass based on wing length ($SBM_{wing}$) as a measure of nestlings' condition (Peig and Green, 2009). This provides an index of wing loading. We used a GLMM with population, female age, and their interaction as fixed factors. Female identity was added as random factor.

We compared clutch initiation date between populations using GLMs with population, female age, and their interaction as fixed factors. Also using GLMs, we compared, clutch size, the duration of incubation, brood size at hatching and the number of nestlings fledged between populations. Population, female age and their interactions were added as fixed factors, and relative clutch initiation day as a covariate. The model for clutch initiation date was fitted with a gamma distribution, and models for clutch size, incubation duration, brood size at hatching and number of nestlings fledged with a Poisson distribution.

GLMs were run using the GENMOD procedure and GLMMs the GLIMMIX procedure in SAS University Edition (SAS Institute Inc., Cary, NC). Post-hoc comparisons were performed using Tukey-Kramer multiple comparison adjustment to obtain corrected p-values. Probability levels <0.05 were considered significant. Data are presented as mean ± SE.

**Results**

*Females' phenotype*

Within each of the four populations, baseline corticosterone level at both the first and third captures was significantly lower than stress-induced and post-dex corticosterone level ($t \geq 4.28$, $p \leq 0.006$; Fig 2). In addition, stress-induced corticosterone level was significantly higher than the post-dex level ($t \geq 3.73$, $p \leq 0.048$; Fig 2).

During mid-incubation, females' baseline and stress-induced corticosterone levels in TN were significantly lower than in WY and AK ($t \geq 4.08$, $p \leq 0.014$; Fig 2). They were also lower in NY than in WY ($t \geq 3.86$, $p \leq 0.031$; Fig 1) but they did not differ between NY and AK ($t \leq 3.00$, $p \geq 0.36$; Fig 2) and between WY and AK ($t \leq 3.13$, $p \geq 0.27$; Fig 3). Post-dex corticosterone level was lower in TN than in WY ($t = 3.96$, $p = 0.022$; Fig 1) but did not differ between NY, WY and AK ($t \leq 3.14$, $p \geq 0.26$; Fig 2).

At the end of incubation, baseline corticosterone was lower in TN than in the 3 other populations ($t \geq 3.67$, $p \leq 0.048$; Fig 2) but did not differ between NY, WY and AK ($t \leq 1.41$, $p \geq 0.98$; Fig 2).

During nestling provisioning, baseline corticosterone levels were lower in TN than NY and AK ($t \geq 5.34$, $p < 0.0001$; Fig 2) but levels in TN did not differ from those WY ($t = 2.69$, $p = 0.61$; Fig 2). In WY, levels were lower than in NY ($t = 4.54$, $p = 0.002$; Fig 2) but did not differ from those in AK ($t = 2.02$, $p = 0.96$; Fig 2). Baseline corticosterone level did not differ between NY and AK ($t = 1.97$, $p = 0.97$; Fig 2). Stress-induced corticosterone levels in TN were significantly lower than in AK ($t = 4.61$, $p = 0.002$; Fig 2) but did not differ from those in NY and WY ($t \leq 1.46$, $p \geq 0.99$; Fig 2). Levels in NY, WY and AK did not differ from one another ($t \leq 2.73$, $p \geq 0.57$; Fig 2). Post-dex corticosterone level did not differ between the 4 populations ($t \leq 1.73$, $p \geq 0.99$; Fig 2).

The corticosterone stress response of females (difference between stress-induced and baseline corticosterone) differed between populations and captures (population x capture number: $F_{3,220.1} = 9.45$, $p < 0.0001$; Fig 3a). During mid-incubation, females in WY had a stronger stress response than females in the 3 other populations ($t \geq 5.08$, $p < 0.0001$; Fig 3a). The stress response was stronger in AK than in TN and NY ($t \geq 2.71$, $p \leq 0.044$; Fig 3a) but did not differ between TN and NY ($t = 1.47$, $p = 0.82$; Fig 3a). During nestling provisioning, females in AK had a stronger stress response than females in the 3 other populations ($t \geq 3.01$, $p \leq 0.048$; Fig 3a) while the response did not differ between TN, NY and WY ($t \leq 0.96$, $p \geq 0.97$; Fig 3a). Within populations, stress response did not change between the two times that it was measured, i.e., during incubation and nestling provisioning ($t \leq 1.49$, $p \geq 0.81$; Fig 3a), except in WY where it decreased from one period to the next ($t = 7.09$, $p < 0.0001$; Fig 3a).

Negative feedback (difference between post-dex and stress-induced corticosterone) differed between populations and captures (population x capture number: $F_{3,236.3} = 8.30$, $p < 0.0001$; Fig 3b). During mid-incubation, females in WY had a stronger negative feedback than females in the 3 other populations ($t \geq 5.36$, $p < 0.0001$; Fig 3b). Females in AK had stronger negative feedback than females in TN ($t = 3.60$, $p = 0.016$; Fig 3b) but similar to that of females in NY ($t = 0.69$, $p = 0.99$; Fig 3b). Negative feedback did not differ between females in TN and NY ($t = 1.55$, $p = 0.77$; Fig 3b). During nestling provisioning, females in AK had a stronger negative feedback than females in the 3 other populations ($t \geq 3.16$, $p \leq 0.04$; Fig 3b) while it did not differ between TN, NY and WY ($t \leq 1.24$, $p \geq 0.91$; Fig 3b). Within populations, negative feedback efficacy did not change between both captures the two times that it was measured ($t \leq 1.28$, $p \geq 0.90$; Fig 3b) except in WY where it decreased between mid-incubation and nestling provisioning ($t = 6.08$, $p < 0.0001$; Fig 3b).

In Tennessee, baseline, stress-induced and post-dex corticosterone did not change between captures ($t \leq 1.68$, $p \geq 0.98$; Fig 2). In New York, baseline corticosterone did not differ between mid and end of incubation ($t = 1.96$, $p = 0.98$; Fig 2) and between end of incubation and nestling provisioning ($t = 2.40$, $p = 0.82$; Fig 2) but was higher during nestling provisioning than during mid-incubation ($t = 4.35$, $p = 0.005$; Fig 2). In Wyoming, baseline corticosterone did not differ between mid and end of incubation ($t = 3.20$, $p = 0.23$; Fig 2) and between end of incubation and nestling provisioning ($t = 0.39$, $p = 1.00$; Fig 2) but was lower during nestling provisioning than during mid-incubation ($t = 3.73$, $p = 0.05$; Fig 2). Stress-induced corticosterone level decreased between incubation and nestling provisioning ($t = 5.26$, $p < 0.0001$; Fig 2) while post-dex level did not change ($t = 1.42$, $p = 0.99$; Fig 2). In Alaska, baseline, stress-induced and post-dex corticosterone did not change between captures ($t \leq 1.71$, $p \geq 0.99$; Fig 2).

Female body mass differed between populations; these differences were affected by capture number (population x capture number: $F_{6,397.5} = 18.73$, $p < 0.0001$; Fig S1). Females in WY were lighter than females in the 3 other populations at all captures ($t \geq 3.73$, $p \leq 0.012$; Fig S1). In TN, females were heavier than in the 3 other populations during mid-incubation ($t \geq 3.56$, $p \leq 0.012$; Fig S1) and their body mass decreased between each capture ($t \geq 8.95$, $p < 0.0001$; Fig S1). In NY, body mass did not change between the first and second captures ($t = 0.16$, $p = 1.00$; Fig S1) but decreased between the second and third captures ($t = 8.09$, $p < 0.0001$; Fig S1). In WY, female body mass did not change captures ($t \leq 1.85$, $p \geq 0.79$; Fig S1). In AK, female body mass decreased between each capture ($t \geq 4.30$, $p \leq 0.0007$; Fig S1).

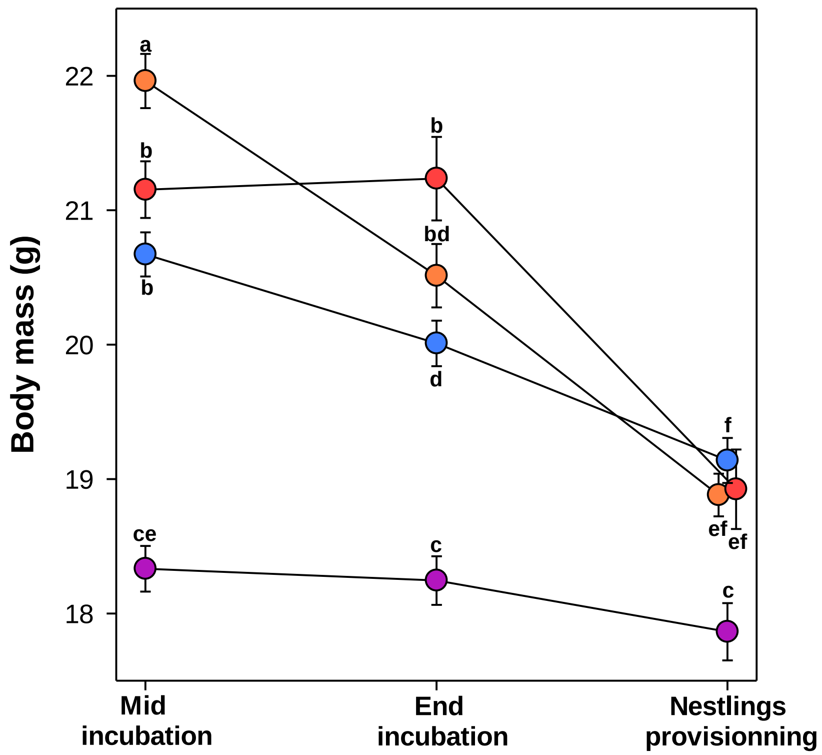


Figure S1: Body mass of females in Tennessee (orange), New York (red), Wyoming (purple), and Alaska (blue) at the first capture during mid incubation, the second capture at the end of incubation and the third capture during nest provisioning. Different letters indicate significant differences.

Glucose level differed between populations and samples (baseline and stress-induced) (population x sample: $F_{3.766.7} = 2.94$, $p = 0.032$). Baseline glucose was lower in WY (196 ± 2 mg.dL$^{-1}$) than in the 3 other populations (TN: 221 ± 2 mg.dL$^{-1}$, NY: 210 ± 3 mg.dL$^{-1}$, AK: 208 ± 2 mg.dL$^{-1}$; $t \geq 3.63$, $p \leq 0.007$) while it was not different between females in TN, NY and AK ($t \leq 0.83$, $p \geq 0.41$). Stress-induced glucose did not differ between the 4 populations (TN: 237 ± 4 mg.dL$^{-1}$, NY: 243 ± 5 mg.dL$^{-1}$, WY: 239 ± 5 mg.dL$^{-1}$, AK: 238 ± 4 mg.dL$^{-1}$;$t \leq 1.66$, $p \geq 0.10$). In the four populations, glucose levels increased significantly between baseline and stress-induced samples ($t \geq 7.67$, $p < 0.0001$).

Resistance to oxidative stress measured by OXY level did not differ between populations ($F_{3,279} = 0.99$, $p = 0.39$) but was higher in SY females (325.44 ± 3.98 mM HOCl) compared to ASY females (306.16 ± 3.77 mM HOCl) ($F_{1,348.1} = 4.24$, $p = 0.040$). OXY level also differed between captures ($F_{2,345.2} = 7.63$, $p = 0.0006$) with lower level during mid incubation (311.98 ± 4.07 mM HOCl) than at the end of incubation (316.17 ± 4.91 mM HOCl) and during nestling provisioning (318.66 ± 5.81 mM HOCl) ($t \geq 2.78$, $p \leq 0.016$) but did not differ between the end of incubation and nestling provisioning ($t = 1.52$, $p = 0.29$). Reactive oxygen metabolites level did not differ between populations and captures (population: $F_{3,285.6} = 0.65$, $p = 0.58$, capture number: $F_{2,274.6} = 2.76$, $p = 0.07$, population x capture number: $F_{6,358.4} = 0.84$, $p = 0.54$).

*Breeding phenology, behavior, nestlings' phenotype and success*

Because tree swallows breeding at higher latitudes arrive later to their breeding sites (Gow et al., 2019), populations differed in their clutch initiation dates (population: $\chi^2_{3,250} = 480.43$, $p < 0.0001$). The four populations differed from each other ($z \geq 3.06$, $p \leq 0.012$) with females in Tennessee (118 ± 1 day of the year) showing the earliest clutch initiation followed by females in New York (135 ± 1 day of the year), females in Wyoming (152 ± 1 day of the year), and finally females in Alaska (158 ± 1 day of the year). Across populations, second-year females initiated clutches later than after-second-year females (relative clutch initiation dates: 14.3 ± 0.8 days vs. 9.0 ± 0.5 days; $\chi^2_{1,250} = 32.63$, $p < 0.0001$).

Clutch size did not differ between populations (TN: 5.8 ± 0.1 eggs, NY: 5.4 ± 0.1 eggs, WY: 5.4 ± 0.1 eggs, AK: 5.2 ± 0.1 eggs; $\chi^2_{3,250} = 5.90$, $p = 0.12$). Clutch size decreased with increasing relative clutch initiation date across populations ($\beta = -0.009$ [-0.018 – -0.0004], $\chi^2_{3,250} = 4.20$, $p = 0.041$) but did not differ with age ($\chi^2_{1,250} = 0.38$, $p = 0.54$).

Incubation duration differed between populations ($\chi^2_{3,159} = 9.01$, $p = 0.029$). Incubation length was the shortest in TN (13.5 ± 0.1 days) that was significantly shorter than in WY where incubation length was the longest (16.4 ± 0.3 days, $z = 2.93$, $p = 0.018$). Incubation duration was intermediate in NY (15.2 ± 0.2 days) and AK (14.5 ± 0.1 days) and did not differ between each other and from TN and WY ($z \leq 2.19$, $p \geq 0.13$).

Brood size at hatching (maximum brood size) did not differ between populations (TN: 4.7 ± 0.2 nestlings, NY: 4.3 ± 0.2 nestlings, WY: 4.4 ± 0.3 nestlings, AK: 4.4 ± 0.2 nestlings; $\chi^2_{3,160} = 1.20$, $p = 0.75$).

Nestling provisioning behavior — measured as the number of daily feeding trips to the nest made by females between day 1 and 1 — increased with increasing brood size ($\beta = 0.041$ [0.035 – 0.047], $F_{1,1933} = 66.13$, $p < 0.0001$). Number of feeding trips by females changed over the nestling rearing period ($F_{17,1933}$

= 654.91, p < 0.0001, Fig. S2). Number of feeding trips generally increased until day 13 when the number of feeding trips reached its maximum before slowly decreasing until day 18 (Fig. S2).

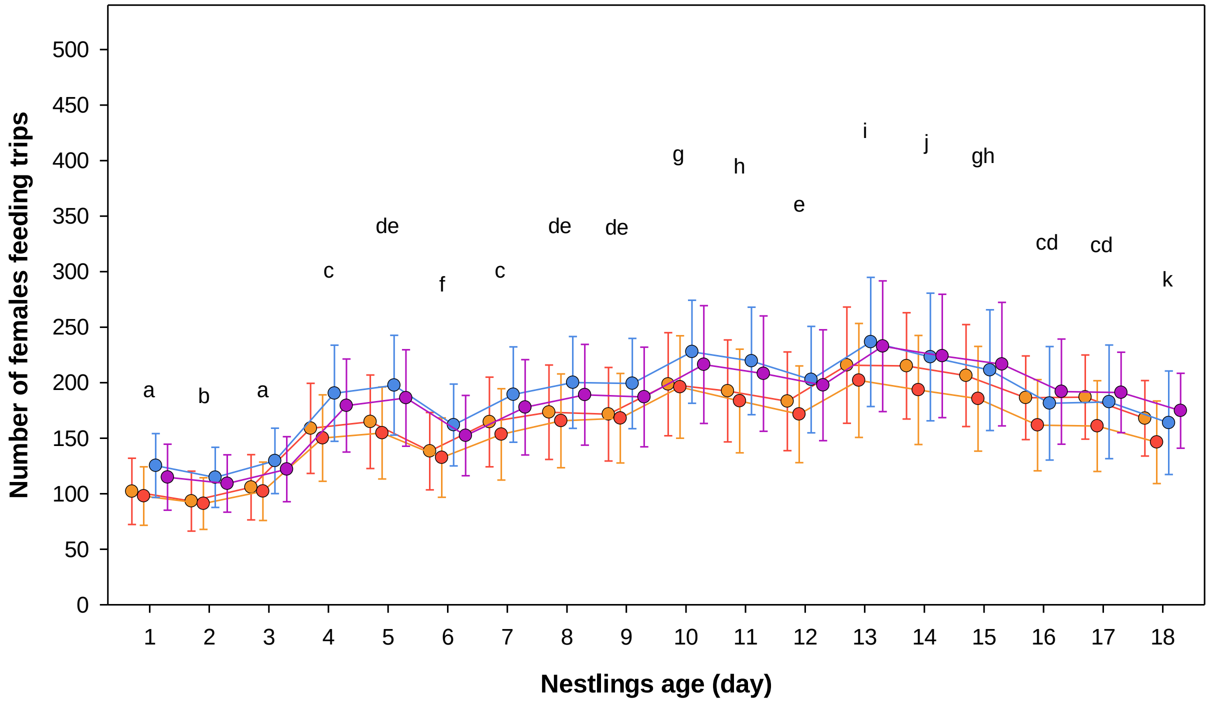


Figure S2: Number of daily feeding trips for females in Tennessee (orange), New York (red), Wyoming (purple), Alaska (blue) over the nestling rearing period. Faded dots represent individual females' data on each day. Bright dots represent average fitted value ± SD for each population for each nestlings' day. Different letters indicate significant differences between days.

Nestling body mass on day 12 after hatching differed between populations ($F_{3,128.5} = 4.02$, p = 0.009). Nestlings in AK (21.2 ± 0.2 g) were significantly heavier than in TN (19.6 ± 0.2 g; t = 2.41, p = 0.018) and NY (18.7 ± 0.5 g; t = 2.65, p = 0.024) but did not differ from those in WY (20.6 ± 0.2 g; t = 1.11, p = 0.68). Fledging success was higher for ASY females (79.2 %, 76 of 96) than for SY females 61.7 %, 50 of 81) ($\chi^2_{1,250} = 5.60$, p = 0.018). Scaled body mass, a measure of condition, was significantly lower in TN than in all other populations ($F_{3,130.5} = 12.06$, p < 0.0001; TN: 18.6 ± 0.2 g, NY: 22.7 ± 0.5 g, WY: 21.2 ± 0.2 g, AK: 21.5 ± 0.2 g; t ≥ 3.01, p ≤ 0.016). The number of nestlings fledged also differed between populations ($\chi^2_{3,250} = 22.49$, p < 0.0001) and age classes ($\chi^2_{1,250} = 4.53$, p = 0.033). The number of nestlings fledged per nest was lower in NY (1.1 ± 0.3 nestlings) than in the 3 other populations (TN: 3.2 ± 0.3, AK: 3.0 ± 0.3,

WY: 2.9 ± 0.3; t ≥ 5.13, p < 0.0001) and greater in ASY (3.1 ± 0.2 nestlings) than in SY females (2.2 ± 0.2 nestlings).

Table S1: Average (±SE) historical hatching and fledging success in the four populations.

| Population | Years | Hatching success | Fledging success |
|---|---|---|---|
| Tennessee | 2014 - 2017 | 81.9 ± 16.6% | 88.6 ± 15.9% |
| New York | 2014 - 2019 | 80.6 ± 3.1% | 78.2 ± 1.9% |
| Wyoming* | 2019 | 95.90% | 91.50% |
| Alaska | 2016-2017 | 70.5 ± 6.2% | 81.2 ± 3.4% |

*Data for Wyoming come from only one year, as this population has not been monitored as extensively as the other populations. Weather conditions during the 2019 breeding season were better than in 2018 with no extended periods of cold and wet weather occurring during the middle of the breeding season. These data show that the Wyoming site is not an inherently poor site for reproduction and hence the low hatching success during the experimental year is likely due to the bad weather conditi0ns during incubation.

**Discussion**

Hatching success and fledging success did not differ much between populations. The exception was that Wyoming females had lower hatching success and New York females had lower fledging success during the experimental year. These two populations experienced unusually bad weather conditions in the year of study during the incubation and nestling rearing periods, respectively. Thus, differences were likely the result of current conditions rather than inherent differences in the quality of these locations as breeding sites (Table S1). In support of this, previous studies at sites in Tennessee, New York and Alaska well as sites in Ontario and North Carolina did not find differences in the number of nestlings fledged (Ardia, 2007; Akçay et al., 2016).

Gow, E. A., Burke, L., Winkler, D. W., Knight, S. M., Bradley, D. W., Clark, R. G., Bélisle, M., Berzins, L. L., Blake, T., Bridge, E. S., Dawson, R. D., Dunn, P. O., Garant, D., Holroyd, G., Horn, A. G., Hussell, D. J. T., Lansdorp, O., Laughlin, A. J., Leonard, M. L., Pelletier, F., Shutler, D., Siefferman, L., Taylor,

C. M., Trefry, H., Vleck, C. M., Vleck, D., Whittingham, L. A. & Norris, D. R. 2019. A range-wide domino effect and resetting of the annual cycle in a migratory songbird. *Proceedings of the Royal Society B: Biological Sciences,* 286**,** 20181916. doi:10.1098/rspb.2018.1916.

Malisch, J. L., Bennett, D. J., Davidson, B. A., Wenker, E. E., Suzich, R. N. & Johnson, E. E. 2018. Stress-Induced Hyperglycemia in White-Throated and White-Crowned Sparrows: A New Technique for Rapid Glucose Measurement in the Field. *Physiological and Biochemical Zoology,* 91**,** 943-949. 10.1086/698536.

Peig, J. & Green, A. J. 2009. New perspectives for estimating body condition from mass/length data: the scaled mass index as an alternative method. *Oikos,* 118**,** 1883-1891. 10.1111/j.1600-0706.2009.17643.x.
